# Diversity and function of maternal HIV-1-specific antibodies at the time of vertical transmission

**DOI:** 10.1101/776856

**Authors:** Laura E. Doepker, Cassandra A. Simonich, Duncan Ralph, Theodore Gobillot, Meghan Garrett, Vladimir Vigdorovich, D. Noah Sather, Ruth Nduati, Frederick A. Matsen, Julie M. Overbaugh

## Abstract

Infants of HIV positive mothers can acquire HIV infection by various routes, but even in the absence of antiviral treatment, the majority of these infants do not become infected. There is evidence that maternal antibodies may provide some protection from infection, but gestational maternal antibodies have not yet been characterized in detail. One of the most studied vertically-infected infants is BG505, as the virus from this infant yielded an Envelope protein that was successfully developed as a stable trimer. Here, we isolated and characterized 39 HIV-specific neutralizing monoclonal antibodies (nAbs) from MG505, the mother of BG505, at a time point just prior to vertical transmission. These nAbs belonged to 21 clonal families, employed a variety of VH genes, many were specific for the HIV-1 Env V3 loop, and this V3 specificity correlated with measurable antibody-dependent cellular cytotoxicity (ADCC) activity. The isolated nAbs did not recapitulate the full breadth of heterologous nor autologous virus neutralization by contemporaneous plasma. Notably, we found that the V3-targeting nAb families neutralized one particular maternal Env variant even though all tested variants had low V3 sequence diversity and were measurably bound by these nAbs. None of the nAbs neutralized the BG505 transmitted virus. Furthermore, the MG505 nAb families were found at relatively low frequencies within the maternal B cell repertoire: all less than 0.25% of total IgG sequences. Our findings demonstrate the diversity of HIV-1 nAbs that exist within a single mother, resulting in a collection of antibody specificities that can shape the transmission bottleneck.

**Importance:** Mother-to-child-transmission of HIV-1 offers a unique setting in which maternal antibodies both within the mother and passively-transferred to the infant are present at the time of viral exposure. Untreated HIV-exposed human infants are infected at a rate of 30-40%, meaning that some infants do not get infected despite continued exposure to virus. Since the potential of HIV-specific immune responses to provide protection against HIV is a central goal of HIV vaccine design, understanding the nature of maternal antibodies may provide insights into immune mechanisms of protection. In this study, we isolated and characterized HIV-specific antibodies from the mother of an infant whose transmitted virus has been well studied.

## Introduction

Mother-to-child transmission of HIV is a unique setting for studying HIV immunity because both the mother and her infant have circulating maternal HIV-specific neutralizing antibodies (nAbs) at the time of HIV exposure and transmission. Antibodies in the mother could potentially neutralize the maternal virus and/or target infected cells to reduce infectiousness. In addition, during late gestation and breastfeeding the infant has HIV-specific antibodies potentially capable of recognizing and blocking maternal viruses through similar mechanisms. However, untreated HIV-exposed infants are still infected at a rate of 30-40%. The specific role of maternal autologous virus-neutralizing IgG responses in driving the selection of infant transmitted founder viruses is both controversial and complex. Some studies, including the larger studies on this topic, report that viruses transmitted from mother to infant are more resistant to neutralization by maternal antibodies than the overall maternal viral population, implying maternal antibodies may select against transmission of the most neutralization sensitive variants (1–3), but this has not been consistently observed in all studies (4, 5). Relatedly, there is also inconsistency in studies that sought to define properties of Env-specific maternal antibodies that are associated with reduced risk of mother-to-child transmission (MTCT) (6). Some studies suggest that protection is associated with antibodies that target specific epitopes such as variable loop region 3 (V3) (7–9) or gp41 (10), while another study found no association between nAb properties and MTCT risk (11) and still others have found antibody specificities that are associated with increased infection risk (12).

Recent studies suggest that passively acquired antibodies that mediate antibody-dependent cellular cytotoxicity (ADCC) provide protection from disease in infants who acquire HIV (13). ADCC antibodies target infected cells for destruction, which have been shown to be a key correlate of MTCT via breastmilk (14). ADCC-mediating HIV specific antibodies in breastmilk have been associated with reduced risk of MTCT in clade A HIV infected women (15), but not in clade C infected women (16). Thus, a deeper understanding of the characteristics, specificities, functions, and potencies of the maternal HIV Env-specific antibody repertoire present at the time of MTCT is warranted.

Given the interactions between the virus and the maternal antibody response, the viruses transmitted to infants are of particular interest. Indeed, one of the most studied HIV-1 variants is from an infant early in infection, BG505 (1), and this infant-derived Env was used to generate the first native-like soluble Env trimer: BG505.SOSIP.664 (17). The BG505 Env has informed a range of structural and immunogen studies (18–21), including a phase I human clinical vaccine trial (https://clinicaltrials.gov/ct2/show/NCT03699241). Infant BG505 was HIV negative at birth but was detected positive at 6 weeks of life, having been infected by a single transmitted variant from mother MG505 (1). At the time of transmission, MG505 had already developed a relatively broad nAb response (22). While the infant was ultimately not protected from the particular transmitted virus that seeded the infection, BG505 did not acquire a myriad of other maternal variants that coexisted at the time of MTCT, as is common in MTCT and HIV infection in general (23). The selection for transmission of a particular maternal variant and not others may have been mediated, in part, by maternal HIV nAbs. In this study, we characterize MG505 Env-specific nAbs that were circulating immediately prior to HIV transmission, focusing on their breadth, potencies, specificities, functions, and frequencies within the antibody repertoire. In addition to measuring nAb efficacy against relevant heterologous viruses, we defined the capacity of these monoclonal maternal nAbs to bind and neutralize seven autologous MG505 viral variants as well as the virus transmitted to infant BG505. Our results demonstrate that there was a diverse repertoire of HIV antibodies in MG505 at the time of transmission, many of which targeted the V3 epitope, bound maternal Envs, and were capable of ADCC.

## Results

### Thirty-nine HIV-neutralizing antibodies were functionally isolated from MG505 immediately prior to MTCT

We isolated and characterized maternal nAbs present very close to the time of vertical transmission of HIV to infant BG505 by studying a maternal peripheral blood mononuclear cell (PBMC) sample from Kenyan subject MG505 collected at 31 weeks of pregnancy (P31). In this case, the P31 time point was 1 week prior to the birth of infant BG505 and 7 weeks prior to the first detection of HIV infected cells in BG505 at six weeks of life (W6). After over 20 years in liquid nitrogen storage, the MG505 PBMCs were 85% viable, containing 5.2 million total live cells. Of live PBMCs, 12% were B cells (CD19+) and, of these, 32% (4% of total cells) were memory B cells (IgM− IgD−). We sorted, cultured, and screened the culture supernatant of 163,165 memory B cells for evidence of antibody activity capable of neutralizing the SF162 Tier 1A HIV-1 variant. This approach identified 190 wells of interest (Figure 1). Monoclonal Abs (mAbs) were successfully reconstructed from 97/190 wells and, ultimately, 39 mAbs were confirmed to neutralize SF162 (Figure 2). Nucleotide somatic hypermutation (SHM) ranged from 4.3-28.6% (heavy chain) and 2.5-10.3% (light chain) and CDR3 lengths ranged from 11-24 (CDRH3) and 9-11 (CDRL3) amino acids (Figure 2). Clonal family analysis revealed that the 39 nAbs comprise 21 clonal families that collectively employ 7 VH genes, 5 VK genes, and 7 VL genes (Figure 2). Two nAbs contained indel events: MG505.18 (family 19) had 6 nucleotides deleted from its VH5-51 FR3 region and MG505.72 (family 8) had 3 nucleotides inserted in its VK3-15 CDR1 region.

**Figure 1.**
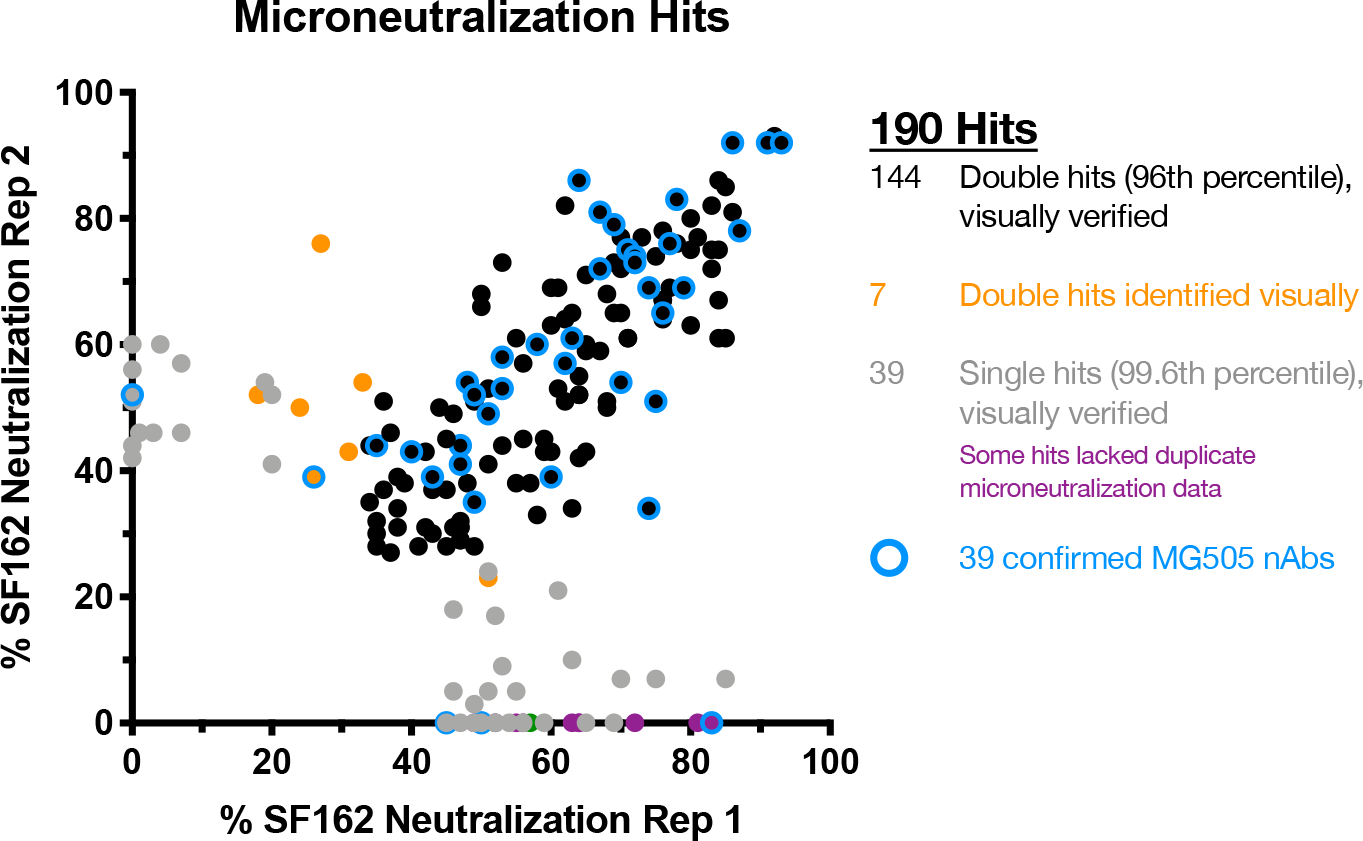
Selection of SF162 pseudovirus-neutralizing MG505 memory B cells for immunoglobulin gene rescue. Wells demonstrating neutralization in the top 96^th^ percentile of wells across all plates for both technical replicates were selected (black) in addition to wells that were in the 99.6^th^ percentile in only one replicate (grey and purple) and seven wells that were subjectively identified by visual inspection as possibly-neutralizing (orange). Wells that were not selected are not shown. Wells that ultimately yielded a confirmed nAb are highlighted in blue.

**Figure 2.**
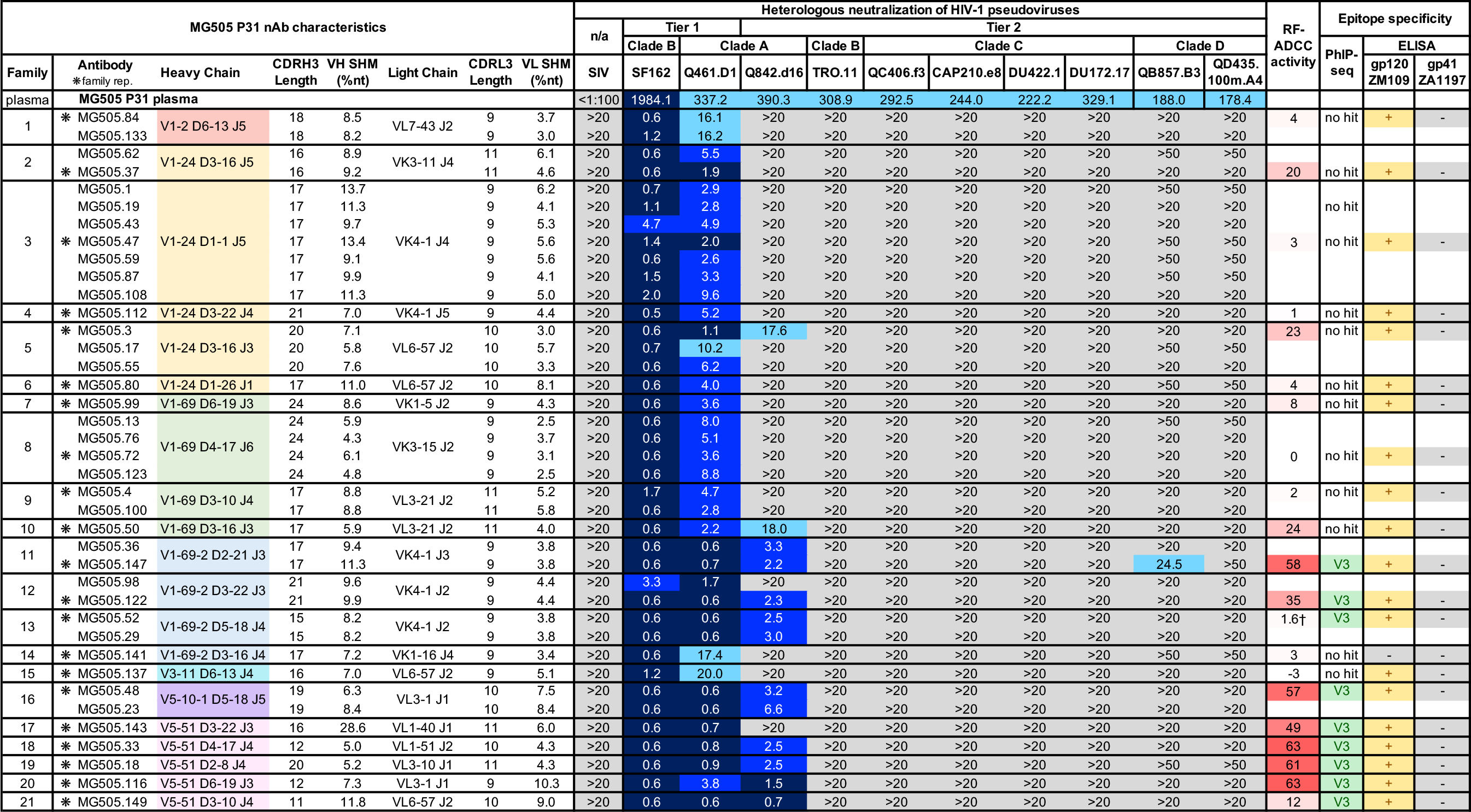
Characteristics of MG505 nAbs. nAb characteristics are displayed in rows, with dark lines separating clonal families. Heavy chain rearrangements are color-coded according to VH gene usage. Neutralization of heterologous viruses by MG505 P31 plasma (top row) and nAbs is displayed as IC_50_ values (reciprocal dilution and μg/ml, respectively). Heterologous virus tiers, clades, and names are indicated. SIV was included as a negative control. Darker blue shading indicates more potent neutralization. Gray indicates that 50% neutralization was not achieved at the highest nAb concentration tested. All nAbs were initially tested at 20 μg/ml, and only if possible low-level neutralization was observed, they were retested at 50 μg/ml. RF-ADCC activity is displayed as percentage of target cells killed, normalized to HIVIG activity, with more potent activity shaded in darker red. PhIP-seq epitope mapping indicates V3 linear peptide specificity for nAb family representatives. ELISA results indicate binding at least 2 times above negative control (+) or lack of binding (−) to indicated antigen. Neutralization, PhIP-seq, and ELISA results reflect averages of at least two independent experiments, each performed in duplicate. RF-ADCC results are representative of two independent experiments. ∗ : nAb selected to represent clonal family, † : RF-ADCC activity varies from 1-27% depending on PBMC donor used in the experiment.

### Tier 1 and Tier 2 heterologous HIV-1 neutralization by MG505 nAbs

To characterize the neutralization potential of the 39 nAbs, we tested each nAb for its ability to neutralize ten heterologous HIV-1 pseudoviruses that were each neutralized by the MG505 P31 plasma (Figure 2). MG505 nAbs all potently neutralized Tier 1 virus SF162 (clade B), which was the major selection criterion for their inclusion in this study, with IC_50_s of 4.7 μg ml^−1^ or lower. They also all neutralized Tier 1 Q461.D1 (clade A), with variable potencies and IC_50_s ranging from 0.6 to 20 μg ml^−1^. A third of the nAbs neutralized a Tier 2 virus from the same virus clade that infected MG505 (clade A, virus Q842.d16) (Figure 2). Only one nAb, MG505.147, demonstrated low potency cross-clade neutralizing activity against Tier 2 virus QB857.B3 (clade D), a variant that is weakly neutralized by MG505 P31 plasma (IC_50_ = 188) (Figure 2). None of the individual nAbs were able to recapitulate the full breadth of the plasma, nor did pooling representatives of the 21 nAb families into a polyclonal mixture (data not shown).

### ADCC functionality correlated with V3 epitope specificity, and Tier 2 heterologous neutralization in MG505 nAbs

Since we previously found that maternal plasma ADCC activity, as measured by the rapid fluorometric ADCC (RF-ADCC) assay, correlated with reduced mortality in HIV-infected infants (13), we further selected one representative nAb from each clonal family to test for RF-ADCC activity against BL035 gp120 (1), a clade A virus isolated from the same cohort as MG505 and BG505. In addition to Tier 1 HIV neutralization, 11 of 21 families were capable of >10% RF-ADCC activity, with values ranging from 12-63%.

To map epitope specificities of each clonal nAb family, we used a combination of phage immunoprecipitation sequencing (PhIP-seq) (24) (Figures 1 and 2) and enzyme-linked immunosorbent assays (ELISA) (Figure 2). For PhIP-seq, we utilized a previously described phage library (24) that contains multiple HIV Env sequences: consensus sequences for clades A, B, C and D and specific sequences circulating in Kenya, including the transmitted BG505.W6.C2 virus. For 9 of the 21 nAb families, phage-displayed peptides were significantly enriched within the V3 region of HIV Envelope (spanning positions 302-322, based on HXB2 numbering), suggesting that this region of HIV Env comprises a key part of the epitope of these isolated nAbs (Figure 3A). Of note, in the case of MG505.149, only a small number of peptides were significantly enriched, which likely led to lengthening of the minimal epitope sequence defined for this nAb through to position 332. Because of this, we have less confidence in the minimal peptide target of this nAb. For three of the nAbs tested (MG505.18, MG505.33, MG505.52), we observed weak but significant enrichments of a peptide that truncated the minimal epitope sequence suggesting this is a core part of the epitope (Figure 3A, blue residues).

**Figure 3.**
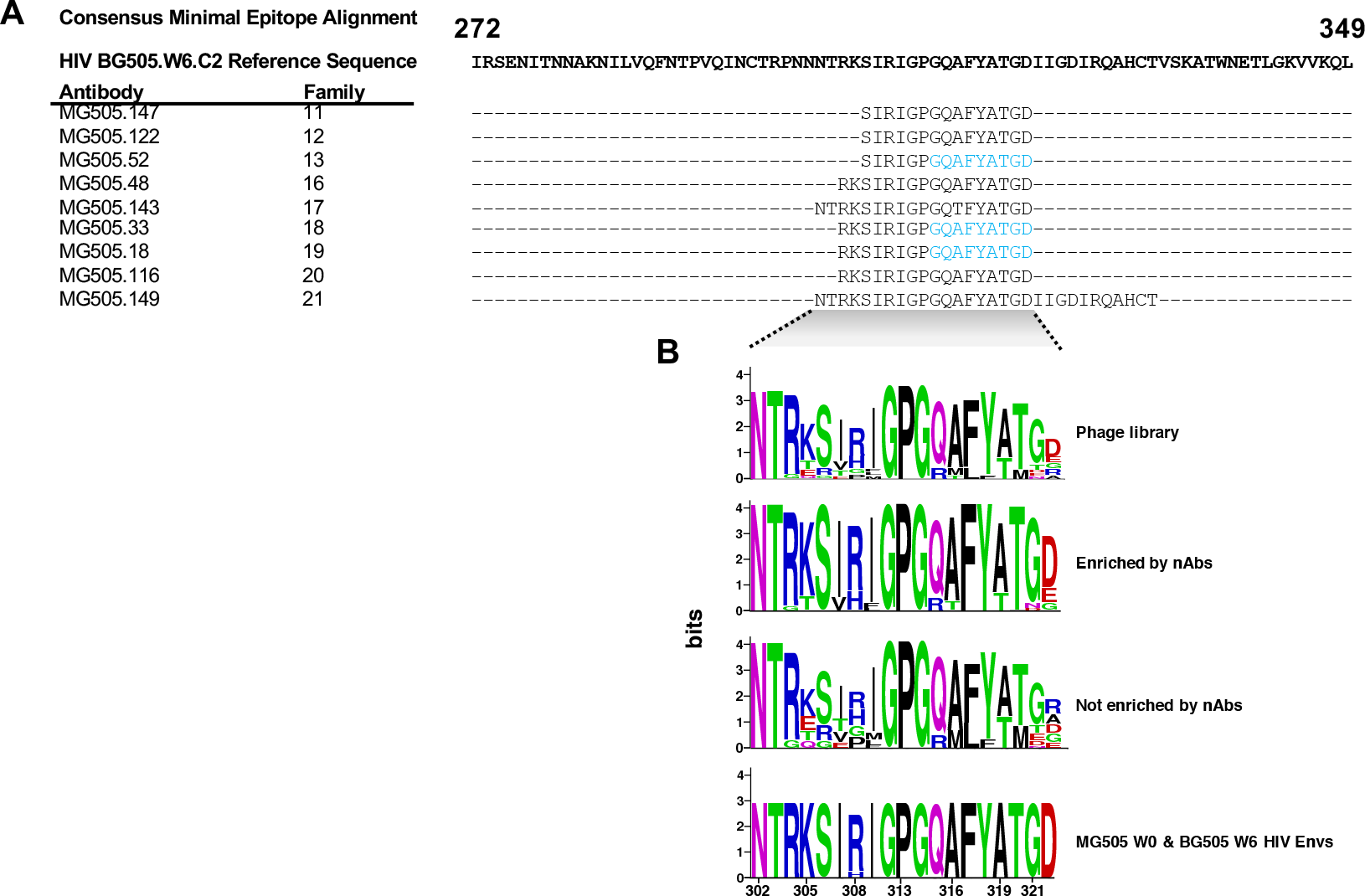
PhIP-seq analysis of nAb family representatives. (A) Sequence alignment of the minimal consensus epitopes identified by PhIP-seq for each tested nAb. See Supplementary Figure 1 for all peptides significantly enriched and not enriched for each tested nAb. Residues in blue signify where the minimal epitope was extended in cases where there was weak but significant enrichment of a peptide that truncated the minimal epitope sequence. (B) Logo plot of sequences corresponding to the minimal epitope region (HIV Env V3) in the phage library, sequences that were significantly enriched or not by tested nAbs, and MG505 W0 & BG505 W6 Envs.

Since the library included sequences of several different HIV-1 variants, our PhIP-seq data also allowed us to gain some insight into which amino acids were preferred at highly variable residues within the library sequences. Interestingly, while we observed some variation in V3 peptide enrichment among different nAbs, there were cases in which peptides with residues at certain positions were consistently enriched while other peptides spanning the same sequence were not enriched. For example, while the phage library contains peptides with amino acids K, T, E, and Q at position 305, the epitope-mapped antibodies only enriched for peptides with K and T at this position (Figure 3B; Supplementary Figure 1). Another example was at positions 321-322, where G and D were enriched, respectively (Figure 3B, Supplementary Figure 1). Of note, in all three cases these preferentially enriched peptides were the same amino acid residues found in the autologous viruses from both MG505 and BG505 Envs (Figure 3B). In all nine cases, the V3-peptides that bound to the nAbs were identical to the BG505.W6.C2 Env V3 sequence (Supplementary Figure 1), which is also identical to the V3 regions of six of seven MG505 W0 Env sequences (Figure 3B and (1)).

These V3-targeting families exclusively employed heavy chain genes VH1-69-2, VH5-10-1, or VH5-51 (Figure 2). There were significant correlations between RF-ADCC activity and V3 specificity (p < 0.0001) and RF-ADCC activity and Tier 2 Env neutralization (p = 0.0015) (Figure 4).

**Figure 4.**
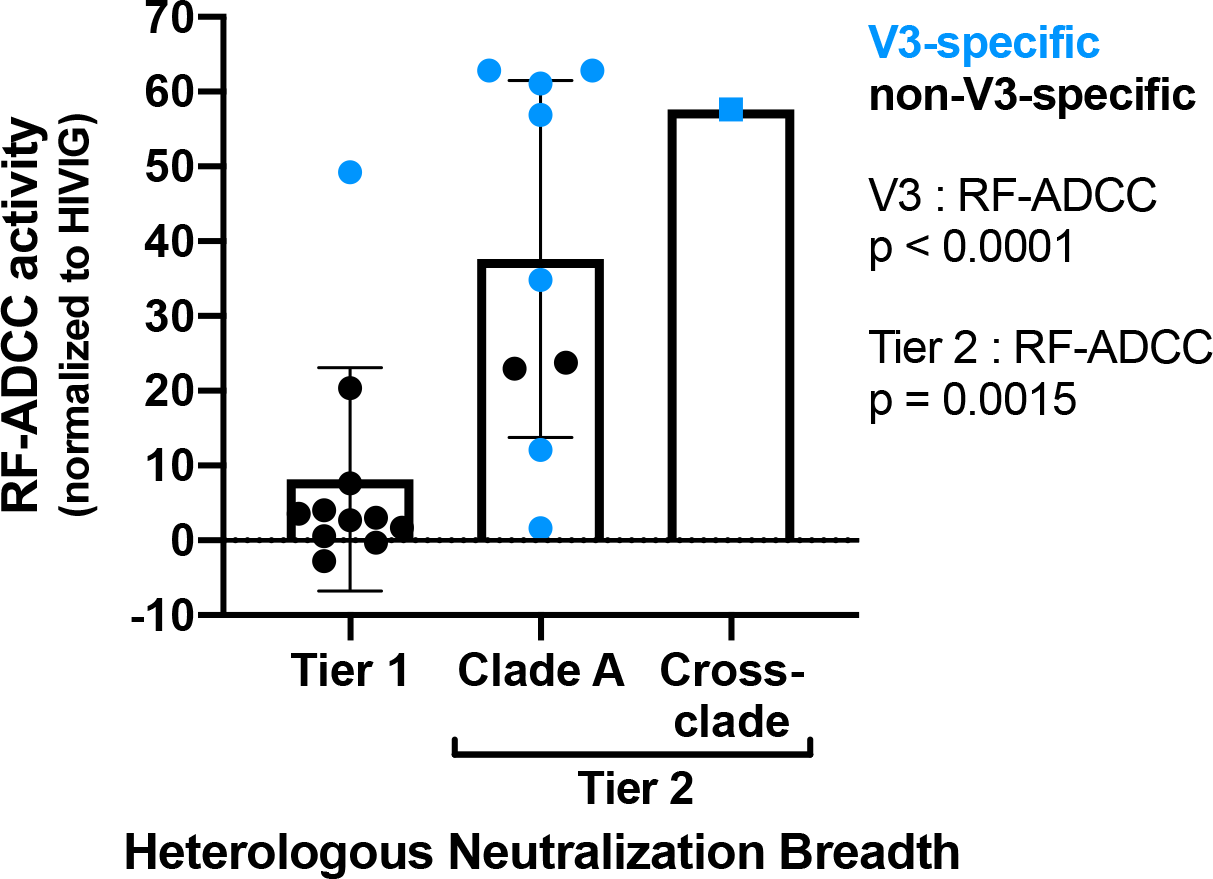
Relationships between nAb functionalities: RF-ADCC activity (y-axis), neutralization (bins), and V3-specificity (blue). Unpaired, two-tailed *t*-tests performed to calculate p values.

The remaining 12 nAb family representatives that were tested by PhIP-seq did not significantly enrich for any phage in the library, suggesting that they may target conformational epitopes that cannot be detected in the 39-mer peptides expressed by phage in the library. To broadly map the epitope specificities of these 12 non-V3-specific nAbs, we employed ELISA assays using gp120 and gp41 antigens. Eleven of twelve families bound gp120 monomer (ZM109) by ELISA; none bound gp41 ectodomain (C.ZA.1197MB) (Figure 2). The single antibody representing family 14, MG505.141, bound neither monomeric form of HIV Env.

### V3-specific nAbs bound, but only rarely neutralized, autologous HIV-1 viruses

Given the relevance of autologous virus neutralization to the prevention of vertical transmission in the context of MTCT, we next tested the nAbs for their abilities to neutralize seven nearly-contemporaneous maternal autologous viruses from the time of birth (W0), which occurred one week after the antibody isolation time point (P31). We also tested three >99.5%-identical infant viruses from the time at which infant HIV infection was first detected (W6). The MG505 P31 plasma neutralized only four autologous viruses, MG505.W0.C2, D1, G2, and H3, and there was no detectable neutralization of the vertically-transmitted viruses tested (Figure 5A). We found that only the V3-specific nAb families neutralized MG505.W0.G2, which was the maternal virus most potently neutralized by the plasma (Figure 5A). The nAbs did not neutralize any other autologous MG505 or vertically-transmitted BG505 viruses.

**Figure 5.**
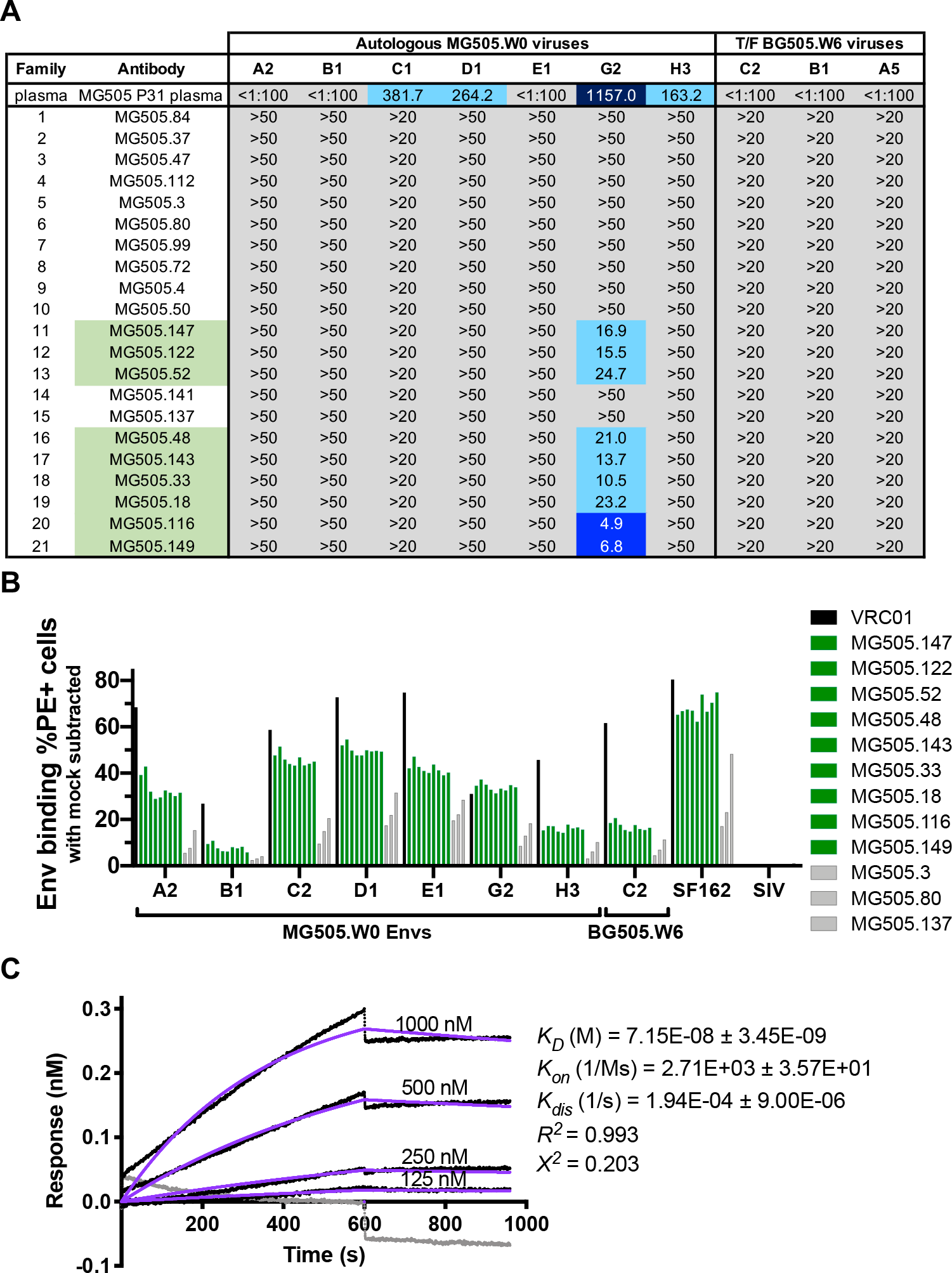
Autologous virus neutralization and binding by MG505 nAbs. (A) Neutralization IC_50_ values of autologous MG505 W0 Envs and vertically-transmitted BG505 W6 Envs by nAb family representatives, reported as averages of at least two independent experiments and displayed for indicated pseudoviruses as in Figure 2. (B) Cell surface autologous Env binding by select nAb families, displayed as mock-subtracted percent of Env-transfected cells bound by each nAb. Data are representative of three independent experiments. Green fill indicates V3-specificity; in gray are nAbs that do not target V3. VRC01 Ab was used as a positive binding control. SIV Env was used as a negative Env control. (C) Biolayer interferometry analysis of MG505.33 mAb (ligand, 8 μg mL^−1^) binding to BG505.SOSIP.664 T332N HIV Env trimer (analyte) at indicated concentrations. The gray line shows 10E8 negative control antibody at 1 μM. Data are representative of two independent experiments. *K*_*D*_, *K*_*on*_, and *K*_*dis*_ are derived from the global best fit (purple) using a 1:1 model of ligand:analyte binding.

The V3 region of MG505.W0.G2 is identical to that of the other maternal and infant viruses, with the exception of MG505.W0.H3 which has a R310H substitution (Figure 3B). To explore the possibility that the V3 nAbs bound the majority of the MG505 and BG505 variants, even though they did not neutralize them, we tested them for Env binding via cell-surface binding assays. Indeed, detectable binding was observed for all V3 nAbs to all autologous Envs expressed on the surface of cells (Figure 5B). We also tested three gp120 nAbs that were not V3-specific. These also bound to cell surface-expressed Env, albeit at lower levels (Figure 5B, shown in gray). Because cell surface-expressed Envs can include various forms of the Envelope protein in addition to the native Env trimer, we more specifically tested whether V3-specific binding, non-neutralizing antibodies could bind the trimeric form of Env. As shown in Figure 5C, the V3-specific MG505.33, which displayed average binding to cell surface-expressed Env, bound to native-like BG505.W6.C2-SOSIP trimer by biolayer interferometry.

### MG505 nAb families were at relatively low frequencies within the maternal B cell repertoire

The relative frequencies of characterized HIV nAb families within a transmitting mother’s larger antibody repertoire are currently unstudied, though these statistics are potentially relevant to understanding which antibody traits permit or prevent MTCT. To better define the frequencies of our 21 nAb families within MG505’s B cell repertoire, we deeply sequenced antibody variable regions (25) from a second P31 MG505 PBMC sample (Table 1). Repertoire clonal family analysis identified IgG sequences clonally related to 3/21 functionally-isolated nAb families: families 3, 9, and 13 (Figure 6A). These nAb families ranked 65^th^, 400^th^, and 278^th^ largest within the IgG repertoire, representing 0.24%, 0.08%, and 0.13% of the total repertoire, respectively. We did not sample any additional clonal IgG sequences stemming from the other 18 nAb families, indicating that family members of these nAbs either absent or rare enough that we did not achieve sufficient sampling depth to detect them (Table 1).

**Table 1.**
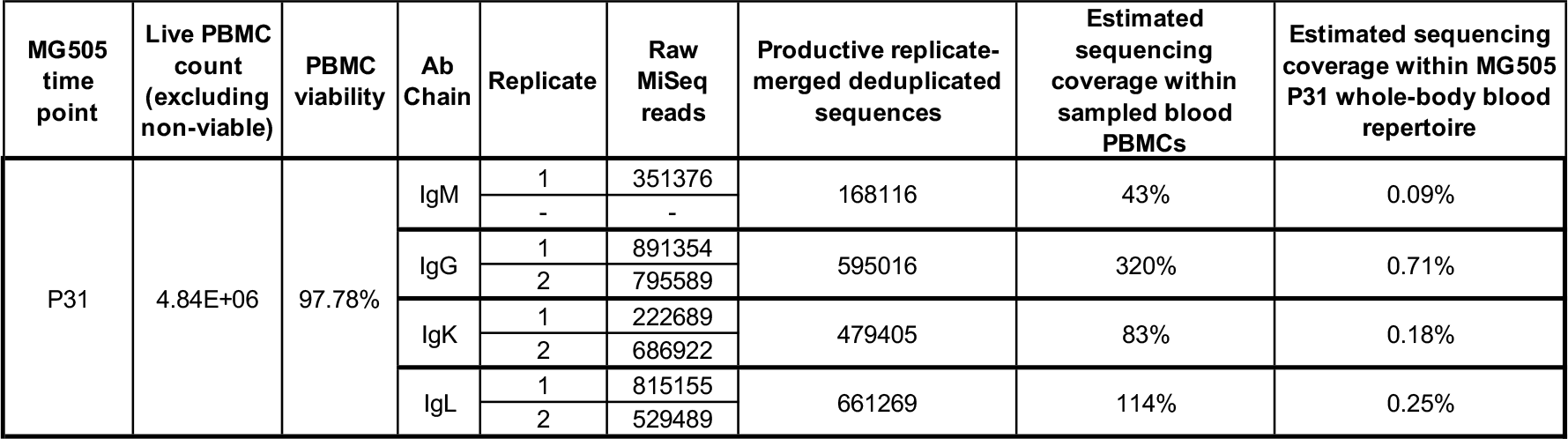
MG505 P31 antibody variable region deep sequencing run statistics. Sequencing coverage was calculated for PBMCs using MG505 P31 B cell frequency statistics from the cell sort of the first aliquot from this time point, as reported in this study. Whole body sequencing coverage was calculated assuming 10 ml of blood was sampled from a total of 4500 ml blood volume.

| MG505 time point | Live PBMC count (excluding non-viable) | PBMC viability | Ab Chain | Replicate | Raw MiSeq reads | Productive replicate-merged deduplicated sequences | Estimated sequencing coverage within sampled blood PBMCs | Estimated sequencing coverage within MG505 P31 whole-body blood repertoire |
| --- | --- | --- | --- | --- | --- | --- | --- | --- |
| P31 | 4.84E+06 | 97.78% | IgM | 1 | 351376 | 168116 | 43% | 0.09% |
|  |  |  |  | - | - |  |  |  |
|  |  |  | IgG | 1 | 891354 | 595016 | 320% | 0.71% |
|  |  |  |  | 2 | 795589 |  |  |  |
|  |  |  | IgK | 1 | 222689 | 479405 | 83% | 0.18% |
|  |  |  |  | 2 | 686922 |  |  |  |
|  |  |  | IgL | 1 | 815155 | 661269 | 114% | 0.25% |
|  |  |  |  | 2 | 529489 |  |  |  |

**Figure 6.**
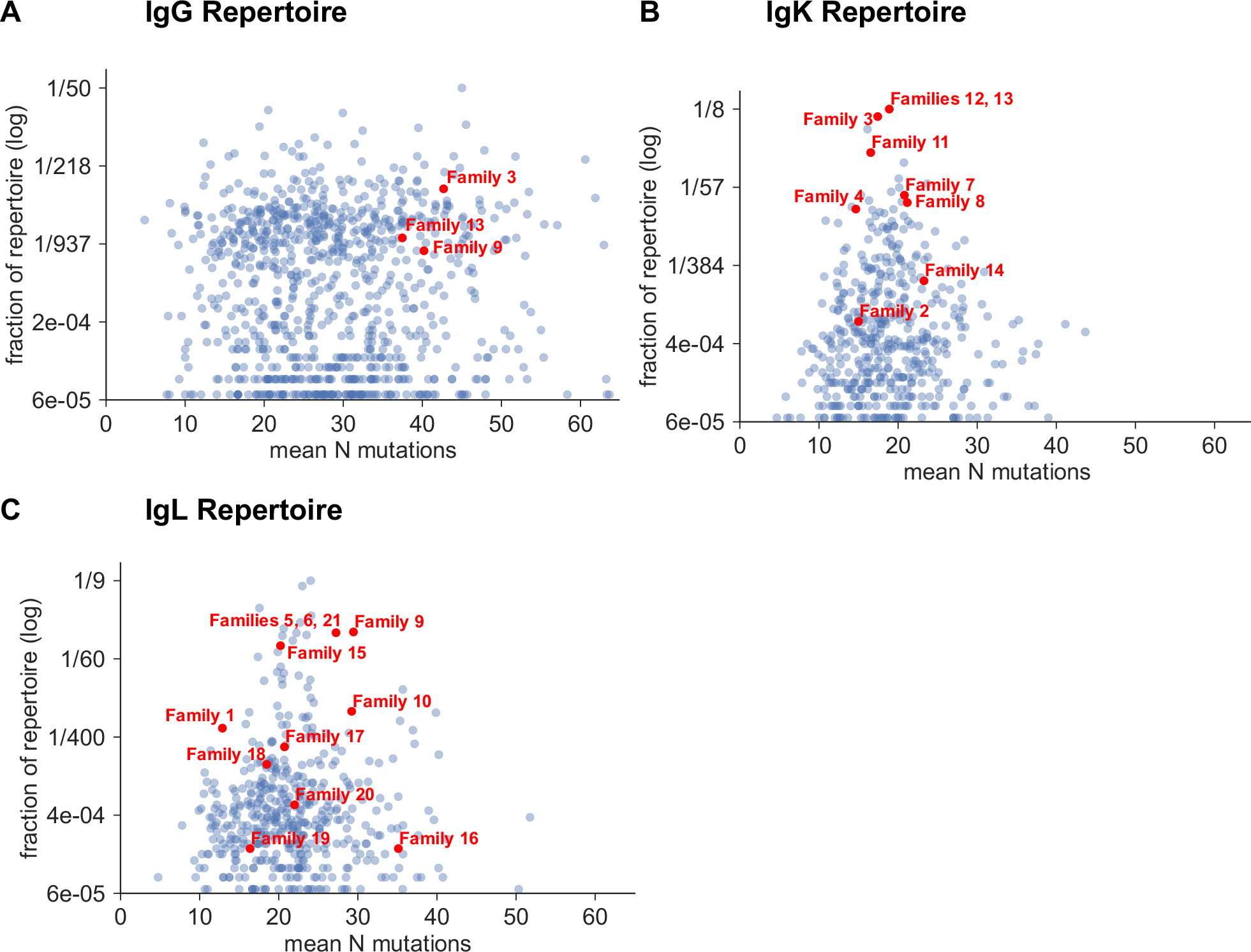
Clonal family analysis of the MG505 P31 B cell repertoire for IgG (A), IgK (B), and IgL (C) antibody variable regions. Each point represents a distinct clonal family, with red points indicating clonal families that contain functionally-identified nAbs. Families smaller than 3 sequences were excluded from the plot.

Repertoire analysis is more nuanced for antibody light chains because of the vastly lower theoretical diversity due to the absence of D genes, shorter CDR3 lengths, and shorter non-templated insertions. The resulting artifactual “superfamilies” reflect examples of preferred light chain usage, where many families with distinct heavy chains have selected the same light chain. Regarding our nAb light chains, we identified IgK or IgL sequences apparently clonally-related to all 21 nAb families, with many families using preferred gene rearrangements that appear as large superfamilies (Figure 6B,C). One example of a superfamily that can be identified unambiguously is the largest family in our kappa analysis, representing 9.85% of the kappa repertoire, which contains members of both nAb families 12 and 13, though we know these are clonally-distinct families based on their paired heavy chains. This effect can also be seen in Figure 6, where the three nAb families found in the heavy chain data appear as much larger fractions of the repertoire in the light chain panels (Figure 6B, C) than in the heavy chain panel (Figure 6A).

## Discussion

Since maternal HIV-targeting antibodies are present at the time of infant infection, MTCT provides a unique setting to explore whether antibodies play a role in determining HIV transmission and infection outcome. This is the first study to examine both the maternal B cell repertoire at the monoclonal antibody level and contemporaneous maternal autologous viruses at the time of MTCT, which, in combination, are clinically relevant to HIV transmission to infants. Here we characterized 39 HIV-neutralizing monoclonal antibodies isolated just prior to vertical HIV transmission in a clade A infected mother. These nAbs were diverse in their relative frequencies, specificities, and functions. Notably, V3-specificity, RF-ADCC activity, and Tier 2 heterologous HIV neutralization were all positively correlated, representing features that may be relevant to evaluating vaccine-induced HIV antibody responses in the future. Of note, while these V3 antibodies neutralized heterologous viruses, most did not neutralize autologous maternal or infant viruses, despite binding to the corresponding infant Env trimer.

The MG505 nAbs were isolated using a high-throughput functional screening method in technical replicate which allowed for less biased discovery of HIV-neutralizing antibodies than Env bait-based approaches, which tend to target antibodies of certain specificities. The diversity of gene usage within the resulting 39 nAbs was striking, especially considering that the screen was not saturating and that there were likely additional families that contributed to the breadth of the MG505 response. Also striking was the fact that we isolated multiple members of 10 clonal families despite their relative rarity in the repertoire, as was indicated by our failure to find additional clonal members of most of these families by deep sequencing. Despite sampling a diverse set of nAbs, including low frequency families, we did not recapitulate the MG505 P31 plasma breadth for heterologous HIV-1 neutralization. Thus, additional nAbs, either from undiscovered families or other clones from families already identified here, likely contributed to the MG505 plasma response. It is possible that 1) the bnAb lineages responsible for the breadth are exceedingly rare populations and we missed them, 2) the antibody-secreting cells (ASCs) and memory cells of the presumed bnAb lineages were not localized to the blood, 3) complex germinal center dynamics resulted in functional differences between the circulating blood memory B cells that we sampled and the plasma antibody-contributing ASCs from the same bnAb lineage, or 4) that the functional screening approach we used did not detect some antibodies. Regardless, the maternal HIV-neutralizing Ab lineages we identified represented <0.25% of the total IgG repertoire. Though relative frequencies of individual nAb lineages are not yet commonly reported, one study reported a comparable value to those we found: adult-derived Protocol C lab code DN HIV-1 bnAb lineage comprised up to 0.8% of the adult’s repertoire at two years post-infection (26). However, it is important to note that antibody lineage frequencies are known to vary greatly across time in humans (27), and that most repertoire sampling, including our own, is limited to blood samples and excludes bone marrow and other important sites of B cell residence.

We used a novel phage immunoprecipitation approach to map the epitopes of these antibodies and found that 9 of the 21 families recognized a linear epitope in V3 that includes the GPGQ sequence that is a common target for V3 nAbs (28, 29). These V3-specific nAbs were capable of both Tier 2 heterologous clade A virus neutralization and ADCC. There is some evidence that V3 nAbs play a role in MTCT, although the results are not consistent across studies and cohorts. V3-specificity was associated with protection in the clade B Women and Infant Transmission Study (7, 8), but not in the clade C Breastfeeding and Nutrition study (11). Based on evidence suggesting that V3-targeting antibodies can select virus escape mutants and drive neutralization resistance in the autologous virus reservoir (30), it has been proposed that maternal V3 nAbs could drive selection of neutralization-resistant transmitted viruses in MTCT (6). Here we found that while the V3 nAbs bound to all autologous viruses tested, they only neutralized one out of seven variants, indicating that the autologous virus reservoir was mostly resistant to maternal V3-specific antibodies present near the time of transmission.

The most potent ADCC-mediating nAbs were all V3-specific. The ability to mediate ADCC was correlated with both V3 specificity and the ability to neutralize a Tier 2 virus. It is possible that V3 specificity could simply enable Tier 2 breadth and ADCC function, especially since V3 antibodies can exhibit weak breadth for Tier 2 viruses due to the viruses’ sampling of the open conformational states that allow for V3 binding (31). Given that ADDC activity has been correlated with infant outcomes (13), these highly potent maternal ADCC nAbs may provide clues to the mechanisms of protection of ADCC antibodies.

It was surprising that the V3 nAbs only neutralized one maternal variant, despite all maternal and infant viruses having identical sequences within the V3 minimal epitope. It is possible that V3 defines only part of the epitope for these nAbs and/or that factors other than minimal epitope binding affect pseudovirus neutralization, such as occlusion of V3 in the native trimer (32). Despite the lack of autologous virus neutralization, we could detect binding to the cell surface-expressed forms of Env of these same viruses. We also detected binding to the BG505 infant Env SOSIP trimer but not neutralization of the corresponding virus. The single difference between the BG505 Envs used in the binding versus neutralization studies is that the native-like SOSIP had an additional glycan at site 332, but this site is outside of the minimal V3 epitope as defined by PhIP-seq. Overall, these studies support the findings of our previous studies of nAbs from infant BF520, which showed that trimer binding and neutralization are not always linked (33).

In the effort to devise an effective HIV-1 vaccine, it is important to understand and target for elicitation protective traits of polyclonal antibody responses that can prevent infection in naïve individuals. Maternal antibodies may provide important insights in the context of MTCT, where only a subset of infants become infected and those that do are often infected with a virus that has escaped maternal antibody pressure. The diverse monoclonal nAb responses described here in the setting of MTCT may provide context for defining the features of antibodies that succeed versus fail at mediating protection against HIV infection.

## Materials and methods

### Human plasma and peripheral blood mononuclear cell samples

Plasma and peripheral blood mononuclear cell (PBMC) samples were from mother MG505 enrolled in the Nairobi Breastfeeding Clinical Trial (34), which was conducted prior to the use of antiretrovirals for the prevention of mother-to-child transmission. The infecting virus was clade A based on envelope sequence (1). Approval to conduct the Nairobi Breastfeeding Clinical Trial was provided by the ethical review committee of the Kenyatta National Hospital Institutional Review Board, and the University of Washington Institutional Review Board.

### B cell sorting

A PBMC sample from MG505 from 31 weeks of pregnancy (P31) was thawed as previously described (33). Cells were stained on ice for 30 minutes using a cocktail of anti-CD19-BV510, anti-IgD-FITC, anti-IgM-FITC, anti-CD3-BV711, anti-CD14-BV711, and anti-CD16-BV711. Cells were then washed once and resuspended in fluorescence-activated cell sorting (FACS) wash (1X PBS, 2% FBS). Cells were loaded onto a BD FACS Aria II cell sorter. The gating strategy was such that memory B cells (CD3− CD14− CD16− CD19+ IgD− IgM−) were sorted into B cell media (IMDM medium, GIBCO; 10% heat-inactivated low IgG FBS, Life Technologies; 5 ml GlutaMAX, Life Technologies; 1 ml MycoZap plus PR, Lonza). Immediately following the sort, memory B cells were plated at 6 B cells in 55 μl per well into 96 × 384-well plates in B cell media supplemented with 100 U ml^−1^ IL-2 (Roche), 50 ng ml^−1^ IL-21 (Invitrogen), and 1×10^5 cells ml^−1^ irradiated 3T3-CD40L feeder cells (ARP 12535). Cultured B cells were incubated for 12 days at 37°C in a 5% CO_2_ incubator based on the protocol by Huang et al. (35).

### B cell culture harvest, microneutralization assay, and reconstruction of antibodies

On day 12, B cell culture supernatants were divided into 2 × 384-well plates at 20 μl each for neutralization assays using a Tecan automated liquid handling system. B cells were frozen at −80C in 20ul RNA storage buffer per well. Microneutralization assays were performed as previously described (33) with one virus in technical replicate (tier 1 clade B SF162). Wells demonstrating neutralization within the 96^th^ percentile in both replicate assays were selected for antibody gene amplification and cloning. Additional wells of interest were subjectively identified by eye, taking into account well position on the plate and surrounding background signal from negative wells. RT-PCR amplification of IgG heavy and light chain variable regions was performed using previously described methods (33). Functional heavy and light chain variable region sequences were determined using IMGT V-QUEST (36). All nAbs were of IgG1 subclass based on 5’ constant region sequencing. Functional variable region sequences were thus cloned into corresponding IgG1, IgK, and IgL expression vectors as previously described (33). In parallel, B cell RNA was sent to Atreca (https://www.atreca.com) for deep sequencing of the antibody heavy and light chain variable regions from each well of interest. In 11 cases where additional heavy and/or light chain sequences were amplified by the Atreca method beyond what we had already identified, the variable chains were synthesized as fragmentGENEs by Genewiz and subsequently cloned into expression vectors. The Freestyle MAX system (Invitrogen) was used to co-transfect paired heavy and light chain plasmids cloned from the same well, and IgG was purified as described (37). For each well, all possible heavy and light chain pairs were generated if more than one antibody was sequenced from the well.

### Pseudovirus production and neutralization assays

Methods for pseudovirus production using envelope-deficient proviral Q23Δ*env* backbone and TZM-bl-based neutralization assays were previously described (38). Plasma IC_50_ values are the reciprocal plasma dilution resulting in 50% reduction of virus infectivity, while monoclonal antibody IC_50_ values represent the mAb concentration (μg/ml) at which 50% of the virus was neutralized. Reported IC_50_ values are an average of at least two independent experiments performed in duplicate. If values from the two experiments disagreed by more than 3-fold, a third experiment was done and all data were averaged.

### ELISA

Immunolon 2HB ELISA plates were coated with 1 μg ml^−1^ ZM109 gp120 monomer or C.ZA.1197MB gp41 ectodomain (Immune Technology Corp.) in 0.1M sodium bicarbonate, pH 9.4 overnight at 4°C. Wells were washed four times with 300 μl wash buffer (1X PBS, 0.05% Tween-20) and blocked for 1 hour at room temperature (RT) in blocking buffer (1X PBS with 10% non-fat milk and 0.05% Tween-20). MG505 nAbs and control antibodies (gp120-specific VRC01 (39) and gp41-specific QA255.006 (24)) were applied at 10 μg ml^−1^ in blocking buffer and incubated at 37°C for 2 hours. Wells were washed and incubated with goat anti-human IgG-HRP (Sigma) diluted 1:2500 in blocking buffer for 1 hour at RT. After washing, TMB-ELISA solution (Pierce) was added for 10 minutes at RT and then stopped with equal volume 1N sulfuric acid. Absorption was read at 450 nM and positive binding was assessed based on blank-subtracted duplicate average absorbance values of at least two times that of the negative control.

### Rapid and fluorometric ADCC (RF-ADCC) assay

The RF-ADCC assay was performed as described (24, 40). In short, CEM-NKr cells (AIDS Research and Reference Reagent Program, NIAID, NIH from Dr. Alexandra Trkola) were double labeled with PKH-26-cell membrane dye (Sigma-Aldrich) and a cytoplasmic-staining dye (Vybrant CFDA SE Cell Tracer Kit, Life Technologies). The double-labeled cells were coated with a clade A gp120 (BL035.W6M.Env.C1, Immune Technology Corp., (1)) for 1 hr at room temperature at a ratio of 1.5 μg protein : 1 × 10^5^ double-stained target cells. Coated targets were washed once with complete RMPI media (Gibco) supplemented with 10% FBS (Gibco), 4.0mM Glutamax (Gibco), and 1% antibiotic-antimycotic (Life Technologies). Monoclonal antibodies were diluted in complete RPMI media to a concentration of 500 ng/mL and mixed with 5 × 10^3^ coated target cells for 10 min at room temperature. PBMCs (peripheral blood mononuclear cells; Bloodworks Northwest) from an HIV-negative donor were then added at a ratio of 50 effector cells per target cell. The coated target cells, antibodies, and effector cells were co-cultured for 4 hr at 37°C then fixed in 1% paraformaldehyde (Affymetrix). Cells were analyzed by flow cytometry (LSR II, BD) and ADCC activity was defined as the percent of PKH-26+ CFDA-cells with background subtracted where background (antibody-mediated killing of uncoated cells) was between 3–5% as analyzed using FlowJo software (Tree Star). All values were normalized to HIVIG (positive control) activity.

### Phage display immunoprecipitation-sequencing

To precisely map the epitopes of antibodies in this study, we employed an approach that combines phage display, immunoprecipitation and highly-multiplexed sequencing, as previously described (24, 41). In brief, amplified phage (1 mL at 2×10^5^-fold representation of each phage clone) that display peptides from several Envelope and full-length HIV sequences was added to each well of a 96-deep-well plate (CoStar). Two concentrations (2 ng, 20 ng) of each monoclonal antibody were then added to phage in technical replicate, with the exception of MG505.52 which was tested at one concentration (20 ng). Phage were subsequently immunoprecipitated and prepared for highly-multiplexed sequencing, as previously described (24).

Bioinformatics analysis of sequencing data was performed using a zero-inflated generalized Poisson significant-enrichment assignment algorithm to generate a −log10(p-value) for enrichment of each phage clone across all samples, as previously described (24). Of note, the −log10(p-value) reproducibility threshold when testing these antibodies in PhIP-Seq was 2.3. Thus, we considered a phage-displayed peptide as significantly enriched if its −log10(p-value) was ≥ 2.3 in both technical replicates. A phage-displayed peptide was considered to be part of the antibody’s epitope sequence only if it was significantly enriched in both conditions tested (2 ng and 20 ng). Fold-enrichment of each phage-displayed peptide was also calculated across all monoclonal antibodies tested.

Phage that were incubated without any monoclonal antibody served as a negative control for non-specific binding of phage and were used to identify and eliminate background hits. For each monoclonal antibody tested, enriched and unenriched peptides were aligned using Clustal Omega. The minimal epitope of an antibody was defined as the shortest amino acid sequence present in all of the enriched peptides. Logo plots were generated using WebLogo (PMID 15173120). For the “phage library” and “not enriched by nAbs” logo plots, only peptides that spanned the full length of the minimal epitope (at least from S308 through D322) were included.

### Analysis of RF-ADCC correlation with heterologous neutralization and V3 specificity

Unpaired, two-tailed *t*-tests were performed using GraphPad Prism 8.

### Cell surface Env binding assays

Binding to cell surface-expressed Env was measured using a flow cytometry-based assay (42). 293T cells (5 × 10^5^ cells) were transfected with indicated 1.33 μg HIV-1 *env* DNA and 2.66 μg Q23gΔ*env* using Fugene6 (Promega), harvested 48 hr post-transfection, and incubated with 20 mg ml^−1^ mAb. Cells were washed and incubated with a 1:100 dilution of goat-anti-human IgG-PE (Jackson ImmunoResearch), washed and fixed with 1% paraformaldehyde, and processed by flow cytometry using a BD FACS-Canto II. Data were analyzed using FlowJo software. Percent binding was calculated as the percentage of PE positive cells with background (mAb binding to cells transfected without *env*, typically 0.2-2%) subtracted. Analyses were performed in GraphPad Prism 8.

### Biolayer Interferometry

MG505 monoclonal antibody binding to HIV Env SOSIP trimer was measured using biolayer interferometry on an Octet RED instrument (ForteBio). Antibodies diluted to 8 μg mL^−1^ in a filtered buffer solution of 1X PBS containing 1% BSA, 0.03% Tween-20, and 0.02% sodium azide were immobilized onto anti-human IgG Fc capture biosensors (AHC). BG505.SOSIP.664 T332N was diluted to 1 μM in the same buffer as above and a series of four, two-fold dilutions of Env trimer were tested as analyte in solution at a shake speed of 600 rpm at 30°C. The kinetics of mAb binding were measured as follows: association was monitored for 10 minutes, dissociation was monitored for 6 minutes, and regeneration was performed in 10mM Glycine HCl (pH 1.5). Binding-affinity constants (K_D_; on-rate, K_on_; off-rate, K_dis_) were calculated using ForteBio’s Data Analysis Software 7.0. Responses (nanometer shift) were calculated using data that were background-subtracted from reference wells and processed by Savitzky-Golay filtering, prior to fitting using a 1:1 model of binding kinetics.

### PBMC RNA isolation for antibody sequencing

PBMCs stored in liquid nitrogen for ~20 years were thawed at 37°C, diluted 10-fold in pre-warmed RPMI and centrifuged for 10 min at 300x*g*. Cells were washed once in phosphate-buffered saline, counted with trypan blue, centrifuged again, and total RNA was extracted from PBMCs using the AllPrep DNA/RNA Mini Kit (Qiagen), according to the manufacturer’s recommended protocol. RNA was stored at −80°C until library preparation. Library preparation, sequence analysis, and antibody lineage reconstruction were performed in technical duplicate, using the same RNA isolated from the MG505 P31 time point.

### Antibody gene variable region sequencing

Antibody sequencing was performed as previously described (25). Briefly, RACE-ready cDNA synthesis was performed using the SMARTer RACE 5’/3’ Kit (Takara Bio USA) using primers with specificity to IgM, IgG, IgK and IgL. cDNA was diluted in Tricine-EDTA according to the manufacturer’s recommended protocol. First-round Ig-encoding sequence amplification (20 cycles) was performed using Q5 High-Fidelity Master Mix (New England BioLabs) and nested gene-specific primers, as previously reported (43). Amplicons were directly used as templates for MiSeq adaption by second-round PCR amplification (20 cycles), purified and analyzed by gel electrophoresis, and indexed using Nextera XT P5 and P7 index sequences for Illumina sequencing according to the manufacturer’s instructions (10 cycles). Gel-purified, indexed libraries were quantitated using the KAPA library quantification kit (Kapa Biosystems) performed on an Applied Biosystems 7500 Fast real-time PCR machine. Libraries were denatured and loaded onto Illumina MiSeq 600-cycle V3 cartridges, according to the manufacturer’s suggested workflow.

### Antibody repertoire sequence analysis and clonal family clustering

Sequences were preprocessed using FLASH, cutadapt, and FASTX-toolkit as previously described (25, 43). Sequences from both technical replicates were combined, deduplicated, and annotated with partis (https://github.com/psathyrella/partis) using default options including per-sample germline inference (44–46). Sequences with internal stop codons, or with out-of-frame CDR3 regions were removed during this step. We did not exclude singletons in an attempt to retain even very rare or undersampled sequences. Sequencing run statistics are detailed in Table 1. Antibody sequences were merged with the 78 functionally-identified MG505 nAb heavy and light chain sequences to form a single comprehensive MG505 P31 antibody sequence dataset. This dataset was then used for clonal family analysis using both the partis unseeded and seeded clustering methods (46). For the unseeded repertoire analysis, since we were interested only in relative properties of clonal families, each data set was subsampled for computational efficiency. For each dataset, three random subsamples of 50,000 sequences were analyzed, comparing results among the three to ensure that they were large enough to minimize statistical uncertainties. No subsampling was necessary for the seeded analysis.

### Neutralizing antibody sequence analysis

Heavy and light chain nAb sequences (“seeds”) were annotated, analyzed, and clustered into clonal families with the MG505 P31 NGS sequences using the partis seeded clustering method on the non-downsampled replicate-merged NGS dataset described above. NAb clusters were delineated for Figure 2 using IgH chain variable region clustering information. Percent SHM was calculated as the mutation frequency at the nucleotide level compared to the predicted naïve allele, as determined by the per subject germline inference for MG505.

### Data availability

The MG505 P31 antibody deep sequencing datasets generated and analyzed during the current study (Figure 6) are publicly available: BioProject SRA accession PRJNA562912 [https://www.ncbi.nlm.nih.gov/bioproject/PRJNA562912]. Maternal nAb sequences reported in this paper have GenBank accession numbers: MN395490-MN395567. The accession numbers for the maternal MG505 W0 and infant BG505 W6 HIV-1 Envs utilized in this paper are GenBank: DQ208449-DQ208455 and DQ208456-DQ208458 (1).

## Acknowledgements

We acknowledge Guy Cavet and Atreca, Inc. for their help with gene amplification and cloning of B cell RNA that resulted in nAb sequences. We thank Brian Oliver, Megan Stumpf, Sonja Danon, Vrasha Chohan, and Mackenzie Shipley for technical assistance with NGS library preparation, antibody production, and neutralization assays. This work was supported by NIH grant R01 AI120961, by training awards T32 AI07140, F30 AI122866, F30 AI142870, and by two new investigator P30 AI027757 awards. The research of F.A.M. was supported in part by a Faculty Scholar grant from the Howard Hughes Medical Institute and the Simons Foundation. The funders had no role in study design, data collection and interpretation, or the decision to submit the work for publication.

---

**Supplementary Figure 1.**
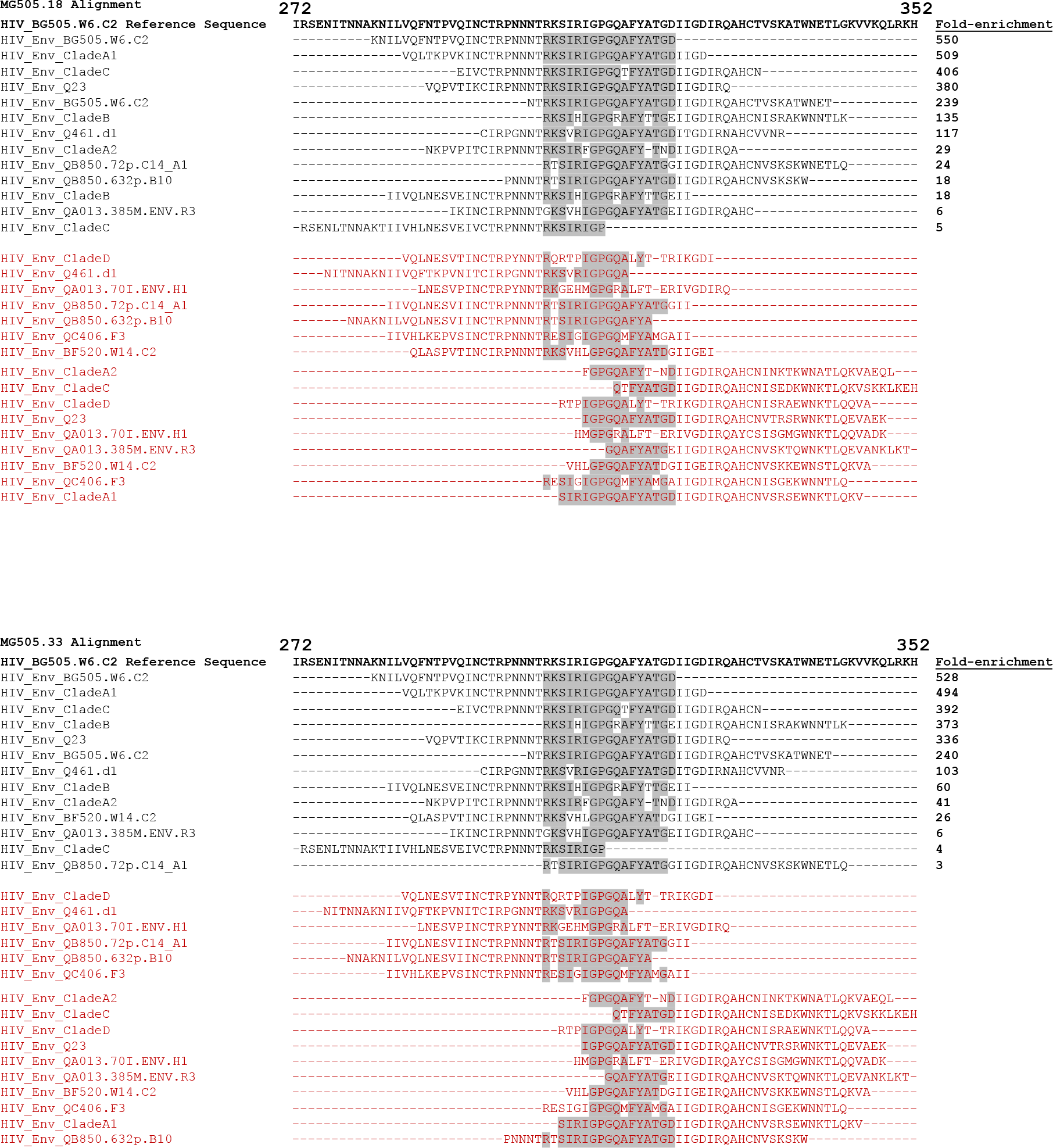
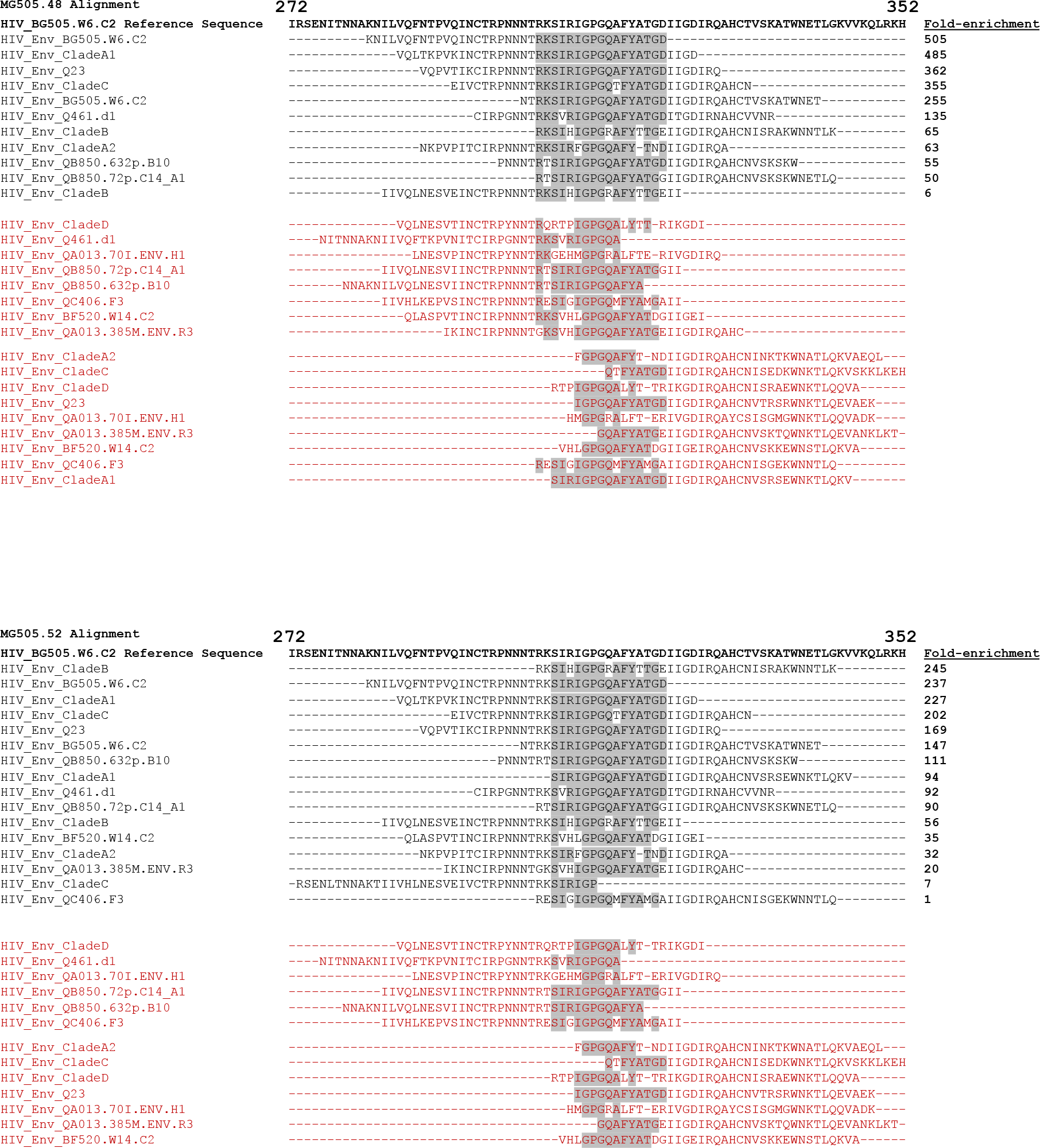
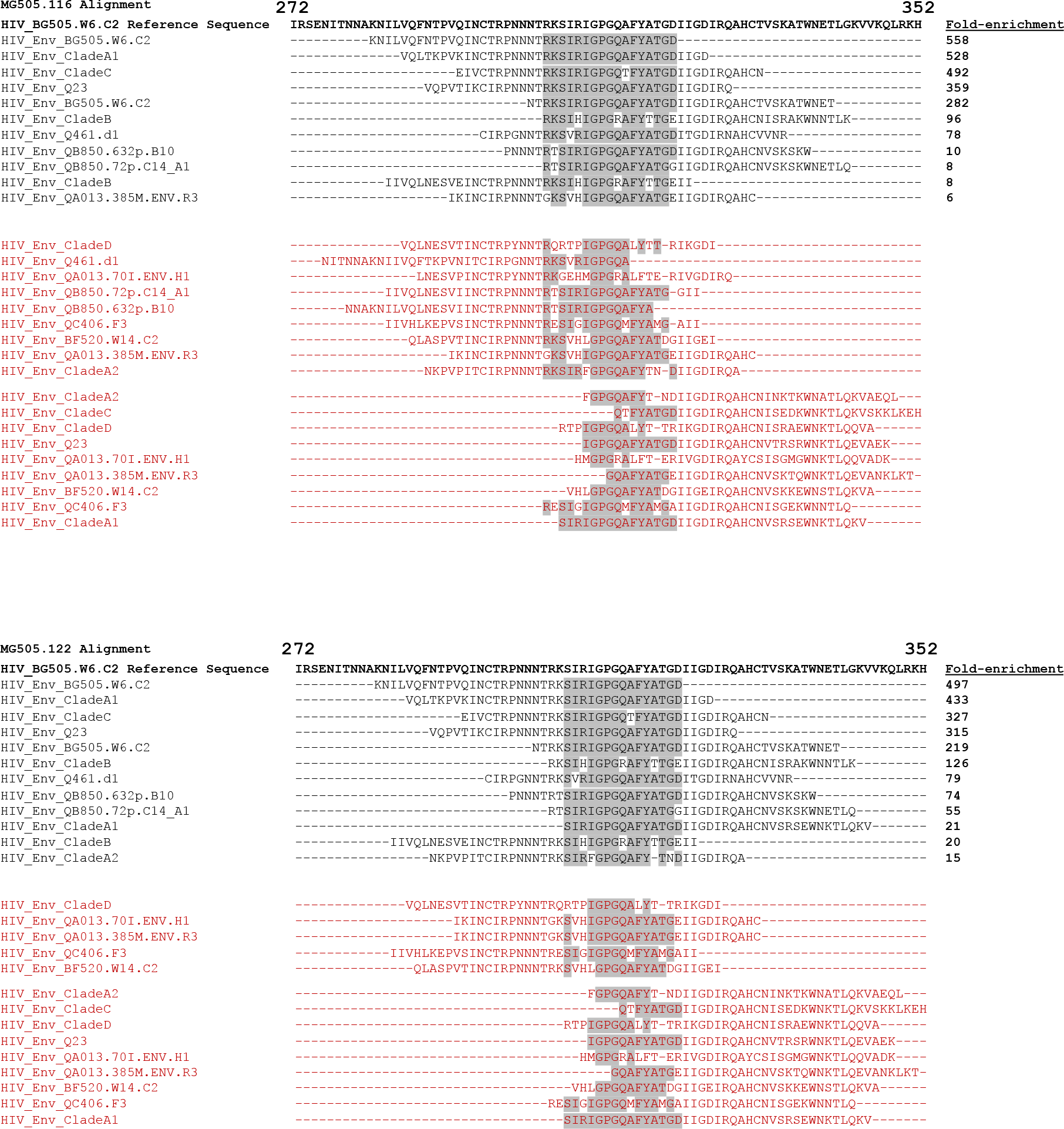
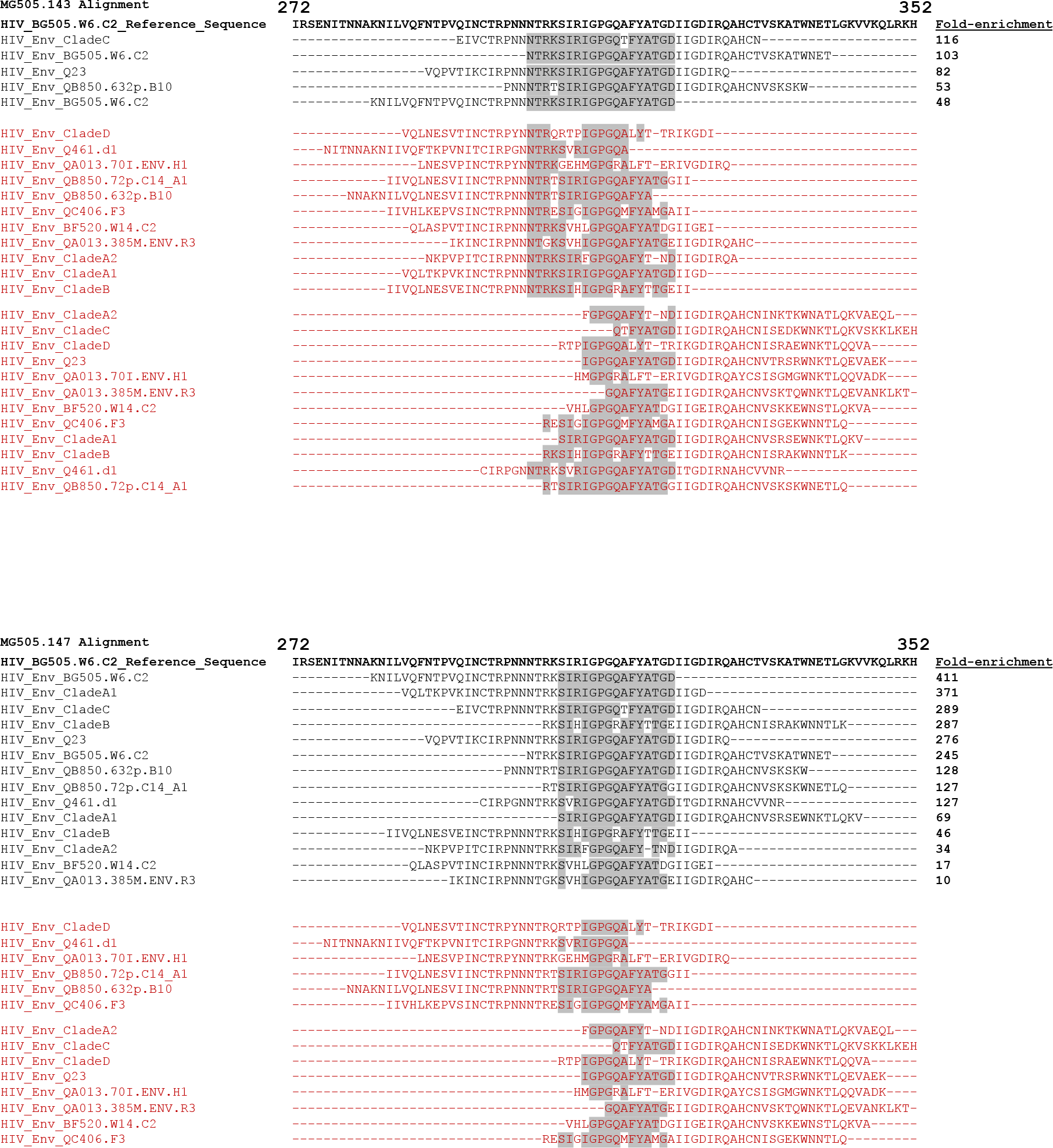
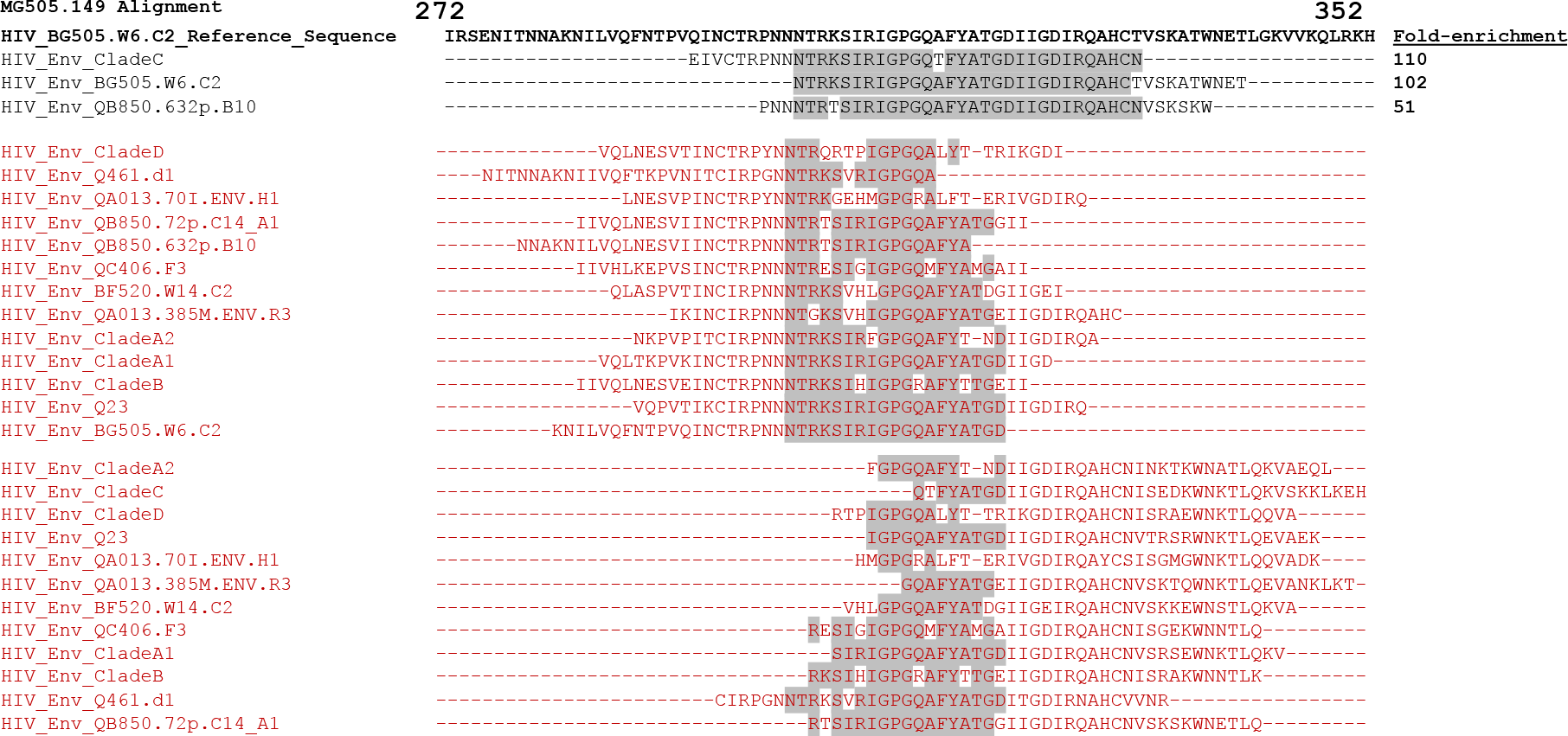
Peptide enrichment for each nAb tested in PhIP-seq. Alignments in black show peptides that were significantly enriched in both tested conditions (2 ng, 20 ng), arranged in descending order of fold-enrichment. Alignments in red show peptides that span this region that were not significantly enriched in both conditions tested. Common sequences among all the enriched peptides are highlighted in gray.

